# Discrete stepping and nonlinear ramping dynamics underlie spiking responses of LIP neurons during decision-making

**DOI:** 10.1101/433458

**Authors:** David M. Zoltowski, Kenneth W. Latimer, Jacob L. Yates, Alexander C. Huk, Jonathan W. Pillow

## Abstract

Neurons in macaque area LIP exhibit gradual ramping in their trial-averaged spike responses during sensory decision-making. However, recent work has sparked debate over whether single-trial LIP spike trains are better described by discrete “stepping” or continuous drift-diffusion (“ramping”) dynamics. Here we address this controversy using powerful model-based analyses of LIP spike responses. We extended latent dynamical models of LIP spike trains to incorporate non-Poisson spiking, baseline firing rates, and various nonlinear relationships between the latent variable and firing rate. Moreover, we used advanced model-comparison methods, including cross-validation and a fully Bayesian information criterion, to evaluate and compare models. These analyses revealed that when non-Poisson spiking was incorporated into existing stepping and ramping models, a majority of neurons remained better described by stepping dynamics, even when conditioning on evidence level or choice. However, an extended ramping model with a non-zero baseline and a compressive output nonlinearity accounted for roughly as many neurons as the stepping model. The latent dynamics inferred under this model exhibited high diffusion variance for many neurons, making them qualitatively different than slowly-evolving continuous dynamics. These findings generalized to alternative tasks, suggesting that a robust fraction of LIP neurons are better described by each model class.

## 1 Introduction

Perceptual decision-making provides an opportunity to probe the role of different brain regions in cognitive tasks (Gold and Shadlen, 2007; Hanks and Summerfield, 2017). In direction discrimination tasks with choices conveyed by a saccadic eye movement (Newsome and Pare, 1988; Britten et al., 1992, 1996), macaque lateral intraparietal area (LIP) responses exhibit positive correlation with choice (Shadlen and Newsome, 1996, 2001). An important series of papers provided support for the idea that the firing rates of LIP neurons reflect the accumulation of sensory evidence in favor of a “preferred” choice target; this hypothesis unified neural responses and behavior under a single theoretical framework known as the drift-diffusion or accumulation-to-bound model (Roitman and Shadlen, 2002; Mazurek et al., 2003; Gold and Shadlen, 2007; Shadlen and Kiani, 2013). An extensive literature has examined this hypothesis in a variety of experimental and theoretical settings (Huk and Shadlen, 2005; Palmer et al., 2005; Ditterich, 2006a,b; Hanks et al., 2006; Kiani et al., 2008; Churchland et al., 2008; Kiani and Shadlen, 2009; de Lafuente et al., 2015)

Although the trial-averaged responses in LIP typically resemble ramps, the average responses do not directly reveal a neuron’s single-trial dynamics. This shortcoming has motivated recent work to determine the single-trial dynamics of LIP responses in direction discrimination tasks (Churchland et al., 2011; Bollimunta et al., 2012; Latimer et al., 2015). In particular, Latimer et al. (2015) compared a discrete switching process or “stepping” model and a accumulation-to-bound or “ramping” model of LIP dynamics, both of which can give rise to ramping trial-averaged activity. They found that the majority of LIP cells were better explained by the stepping model. However, subsequent literature has sparked debate over the interpretation of these results (Shadlen et al., 2016; Zylberberg and Shadlen, 2016; Chandrasekaran et al., 2016; Latimer et al., 2017; Zhao and Kording, 2018).

In this paper, we aim to settle the debate about single-trial LIP dynamics using improved models and model comparison methods. We have extended the classic ramping and stepping models of LIP dynamics in several important ways. First, we incorporated spike-history dependencies into both models to account for the temporal autocorrelations of spike trains in LIP. Second, we investigated nonlinear ramping models with a non-zero baseline firing rate and several possible nonlinear relationships between the latent variable and firing rate. We compared these models using a principled, fully Bayesian information criterion and Bayesian leave-one-out cross-validation.

In our analyses, the stepping model outperformed the ramping model for a majority of neurons when both models were extended to incorporate spike-history. This result was robust to partitioning of the data by choice or sensory evidence level, showing that (in contrast to recent analyses in Zylberberg and Shadlen (2016)) model selection was not driven by anti-preferred choice or evidence trials. On the other hand, an extended ramping model with a non-zero baseline firing rate and a decelerating nonlinearity outperformed the stepping model for slightly more than half the neurons in our population. The latent firing rates inferred under this model, however, were often governed by high diffusion variability, making the distinction between continuous and discrete models less concrete. These analyses revealed that spike responses in LIP are more complex than simple ramping or stepping models, while confirming that discrete dynamics provide the best account for a substantial fraction of neurons in LIP.

## 2 Results

We formulated explicit statistical models of latent dynamics underlying single-trial spike trains in area LIP during perceptual decision-making and used two different statistical methods to compare them. Our analysis builds on Latimer et al. (2015), which formulated the ramping and stepping latent variable models of LIP spike trains. The basic ramping model, often referred to as the drift-diffusion or accumulation-to-bound model, consists of a continuous latent diffusion process, which is passed through a soft-rectifying nonlinearity to obtain a Poisson firing rate. The basic stepping model, on the other hand, consists of a discrete switching process that jumps from an initial firing rate to one of two levels with a probability that depends on the stimulus. Here we extended these two models in order to incorporate non-Poisson spike-history effects and to allow additional forms of nonlinearity in the ramping model (Figure 1).

**Figure 1.**
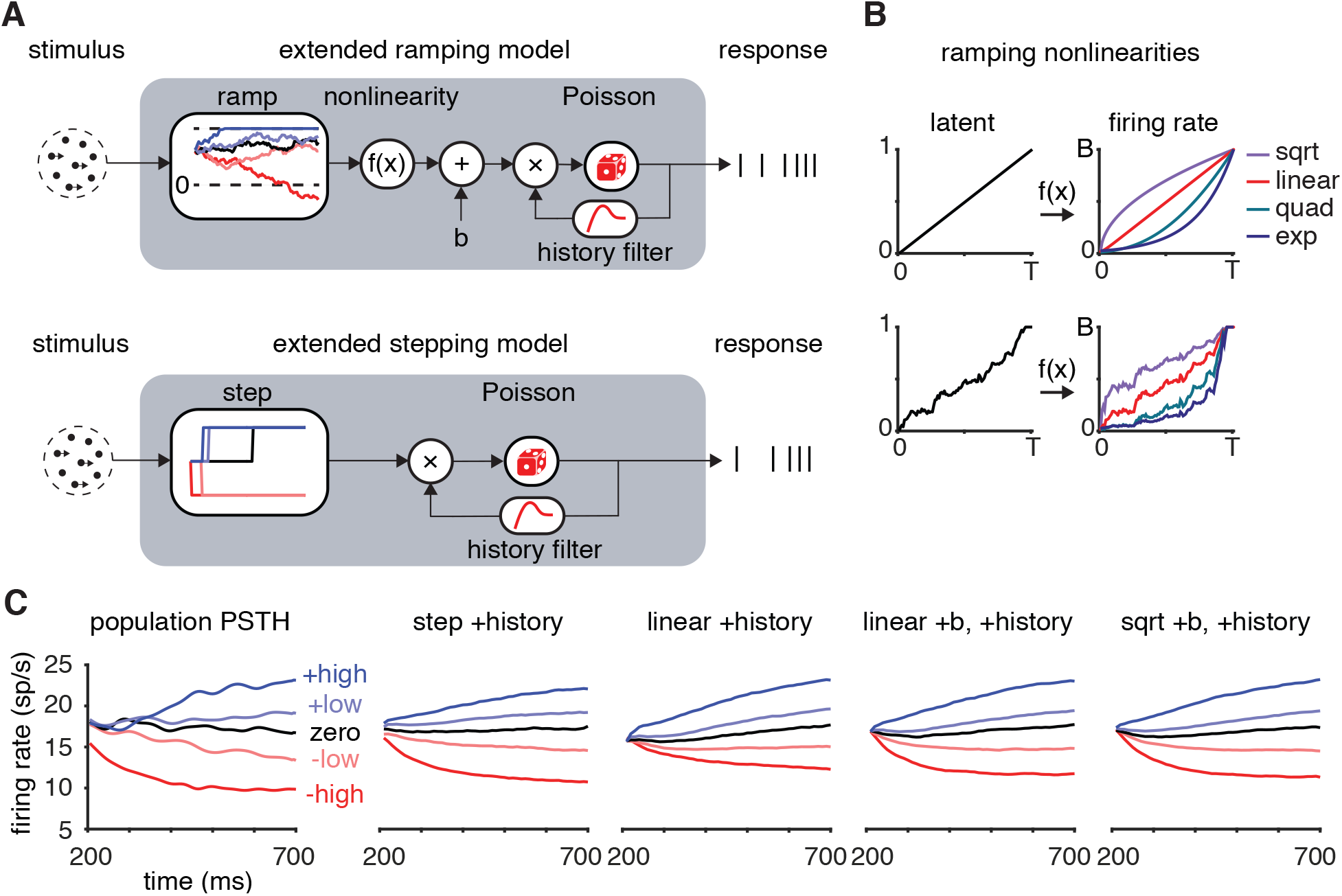
Latent variable models for LIP spike responses during decision-making. **A**. Schematics of extended ramping (above) and stepping (below) models. For the ramping model, the stimulus coherence sets the drift rate of a latent diffusion-to-bound process; this process is transformed by a rectifying nonlinearity *f*(·) and added to a baseline firing rate *b*. For the stepping model, stimulus coherence determines the distribution over time and direction of a discrete step to one of two possible latent firing rates. In both models, the latent firing rate is multiplied by the exponentiated output of a spike history filter, allowing it to capture non-Poisson firing statistics. **B**. Output nonlinearities considered for the extended ramping model. The nonlinearity transforms example latent diffusion paths (left) into different latent firing rate trajectories (right). **C**. Average firing rates for different motion coherence levels, aligned to motion onset and averaged across neurons (left), along with averaged responses simulated from models considered in this paper, where “+b” indicates inclusion of a non-zero baseline and “linear” and “sqrt” indicate the choice of nonlinearity in the extended ramping model.

To compare these models, we used two different methods: the Watanabe-Akaike information criterion (WAIC, Section 4; Watanabe, 2010; Gelman et al., 2014) and Bayesian leave-one-out cross-validation (Vehtari et al., 2017). The WAIC has multiple features that make it robust for model comparison. First, it uses the full posterior over the parameters for model evaluation, and therefore does not rely on a point estimate of the parameters (which is the case for other model-selection criteria, e.g., AIC, BIC, or DIC). Also, the penalty term in the WAIC is stable and guaranteed to be non-negative, in contrast with the DIC (Gelman et al., 2014; Vehtari et al., 2017). Finally, the WAIC has solid theoretical grounding as it is asymptotically equivalent to Bayesian leave-one-out cross-validation (Gelman et al., 2014; Vehtari et al., 2017). We find these benefits are realized empirically, as the WAIC outperforms the DIC at identifying the true model in simulations (Figure 7B).

### 2.1 Incorporating spike-history dependencies

The basic ramping and stepping models from Latimer et al. (2015) described spiking as Poisson conditioned on the latent ramping or stepping process, which ignores spike-history effects present in real spike trains (e.g. refractoriness, bursting, spike-rate adaptation). We therefore extended both models to include autoregressive spike-history filters, like those in the generalized linear modeling (GLM) framework (Figure 1, Section 4). These filters capture non-Poisson spike-history dependencies (Truccolo et al., 2005; Weber and Pillow, 2017) and allow for a dissociation of latent dynamics from spiking activity that can be explained more parsimoniously by past spikes.

We fit stepping and ramping models both with and without spike-history to the responses of 40 LIP cells during a variable-duration random dot motion task (Section 4; Meister et al., 2013). Simulated data from the fitted models captured the shape of true population PSTHs for different coherence levels (Figure 1C). The inferred spike history filters typically exhibited short-timescale inhibition (or refractoriness) and longer-timescale self-excitation, although there was substantial variability across neurons (Figure 2A). These filters conferred a dramatic improvement in the ability to capture temporal auto-correlations in spiking activity under both models (Figure 2B). Intriguingly, the history filters were nearly identical between stepping and ramping models for the majority of neurons. This suggests that the filters captured similar structure across models, and were not strongly influenced by model-specific assumptions about latent dynamics. Spike-history filters substantially improved prediction accuracy under both models for the vast majority of neurons, giving better WAIC for 38 out of 40 cells (Figure 2C).

**Figure 2.**
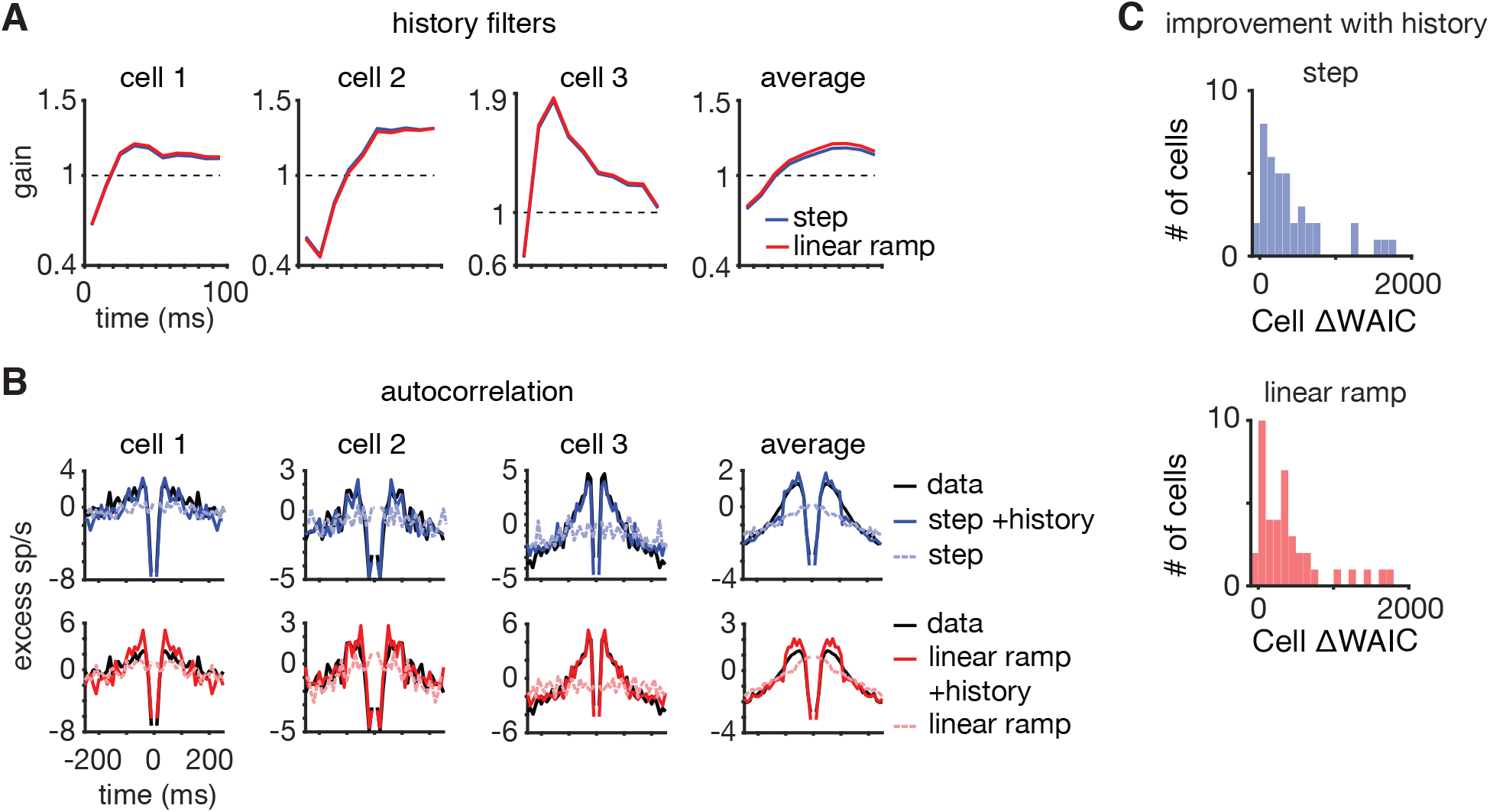
Models with spike-history dependence outperformed classic stepping and ramping models. **A.** Spike-history filters for three example neurons, and the average spike-history filter across neurons, for extended stepping (blue) and extended ramping (red) models. Inferred filters were remarkably similar between models, implying that spike-history effects did not vary with the choice of latent dynamics model. **B**. Spike train autocorrelations of LIP neurons and models with and without spike-history filters, revealing that classic stepping and ramping models could not account for the temporal statistics of real spike trains. **C**. Models with spike-history filters performed better than classic stepping (above) and ramping models (below) for the majority of LIP neurons, as quantified by WAIC. Positive WAIC differences favor the model with spike-history.

### 2.2 Stepping model robustly outperforms linear ramping model

Model comparison using the WAIC and cross-validation revealed that the stepping model with spike-history outperformed the linear ramping model with spike-history for 28 out of 40 cells (Figures 3A and 8A). We quantified uncertainty in the model comparison using the standard error of the WAIC difference across trials (Section 4). For the 26 cells whose WAIC differences were more than the standard error from zero, 21 favored the stepping model. This result was not driven by the penalty term in the WAIC, as the stepping model had higher predictive accuracy for 33 of the 40 cells (higher lppd, Section 4, Equation 68).

**Figure 3.**
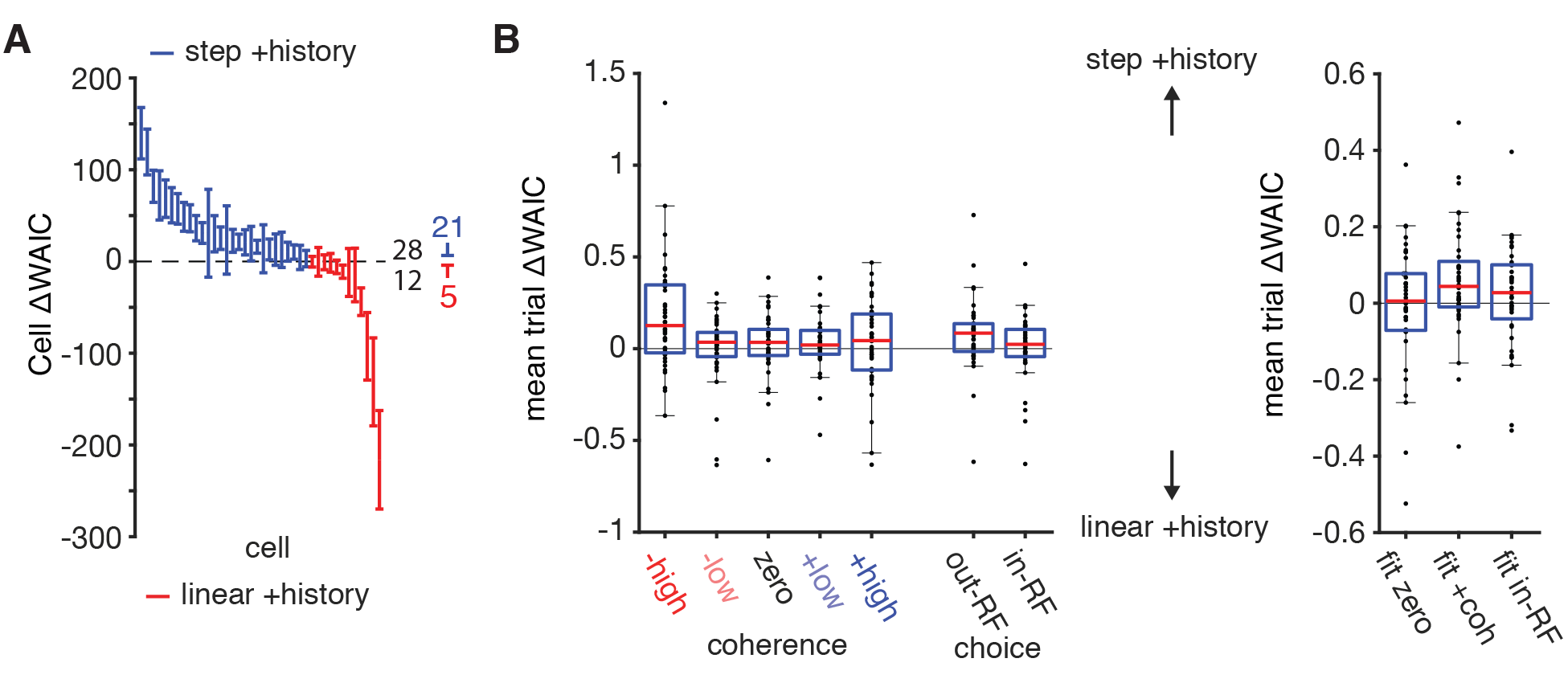
Stepping model with spike-history outperforms linear ramping model with spike-history for a majority of cells across conditions. **A**. Sorted WAIC differences between the stepping and linear ramping models with spike-history for all neurons (error bars indicate *±*1SEM). Blue (red) lines indicate cells for which the stepping (ramping) model had a better WAIC value. Colored numbers indicate the number of cells where the WAIC difference favored stepping or ramping by more than the standard error. **B**. *Left*: WAIC differences computed for subsets of trials conditioned on motion coherence and choice. The differences were normalized by the number of trials in each condition. Positive differences favor the stepping model. *Right*: WAIC differences for models fit only to data from zero coherence, positive coherence, or in-RF trials (respectively), normalized by the number of trials in each condition. Similar results were obtained with cross-validation (Figure 8).

To examine how different trial conditions contributed to the model comparison across cells, we then compared the models across subsets of trials partitioned by the motion coherence and choice. For each coherence and choice, the stepping model with spike-history outperformed the linear ramping model with spike-history for the majority of cells (Figures 3B and 8B). The negative high coherence trials had the largest median difference between cells, which reflects concerns about the inability of the ramping model to handle strong negative drifts in firing rate (Shadlen et al., 2016; Zylberberg and Shadlen, 2016). However, the median difference favored stepping for each coherence level, including those for weak or strong motion into the RF. Also, with the negative high coherence or all negative coherence trials excluded, the stepping model still outperformed the ramping model for a majority of cells (26 out of 40). This suggests that the overall comparison did not depend on the negative coherence trials and that the dynamics were consistent across stimulus conditions and choice.

We next evaluated the possibility that inference of the model parameters was influenced by subsets of trials, which would also affect the model comparison. We re-fit the models to three different subsets of trials (Figures 3C and 8B). The first contained all zero-coherence trials, which putatively have long integration times (and therefore might be expected to have the slowest or most gradual “ramp-like” dynamics). The other two re-fits used data from all positive coherence trials and data from all in-RF choice trials, in which the animal made a saccade to the “preferred” target. These latter two fits restricted analysis to trials with putatively positive values of accumulated sensory evidence. For all three analyses, model comparison favored the stepping model with spike-history for more than half the neurons: 21/40 for zero-coherence trials, 25/40 for positive coherence trials, and 27/40 for in-RF choice trials.

Recent work has argued that trials with strong initial negative diffusion and termination at non-zero rates might bias model comparison in favor of the stepping model (Zylberberg and Shadlen, 2016). The competing accumulators model assumes that these trials occur most prevalently during negative coherence trials or out-RF choice trials (Mazurek et al., 2003). These are trials where either the presented or perceived evidence is to the target outside the RF of the LIP neuron, and therefore are trials where the LIP neurons are more likely to decrease their firing rates throughout a trial. Given that assumption and the analyses performed above, we conclude that the comparison between the ramping and stepping models with history fit to all trials was not driven by these trials, even though a mechanism such as a baseline firing rate to stop strongly going negative rates was not included.

### 2.3 Nonlinearities and non-zero baselines improve the ramping model

Although the ramping model formulated in (Latimer et al., 2015, 2017) was motivated to capture the hypothesized linear relationship between a biased diffusion-to-bound process and single-neuron firing rates, neurons might exhibit nonlinear relationships between a putative latent diffusion process and their firing rates. To investigate this possibility, we fit nonlinear ramping models with a variety of different nonlinearities: a soft-rectified square root function (“sqrt”), a soft-rectified quadratic (“quad”), or an exponential (“exp”) (Section 4). These models can capture varying degrees of nonlinear response saturation or acceleration as a function of the latent variable (Figure 1B). We also included a non-zero baseline firing rate parameter *b* that acts as a (non-absorbing) lower bound on the firing rate (Section 4). This prevents firing rate from going to zero when the drift term is strongly negative (Shadlen et al., 2016; Zylberberg and Shadlen, 2016; Latimer et al., 2017).

For each nonlinearity, we fit the ramping model with and without an additive baseline and spike-history filters. We compared each model to the original linear ramping model without spike-history from Latimer et al. (2015) (Figure 4A). Including the non-zero baseline improved all models, and models with the largest median improvements over the linear ramping model included both the non-zero baseline and spike-history filters. Across all nonlinear ramping models with baseline, the model with square root nonlinearity performed best, achieving the best WAIC for more than 50% of neurons (Figure 4B).

**Figure 4.**
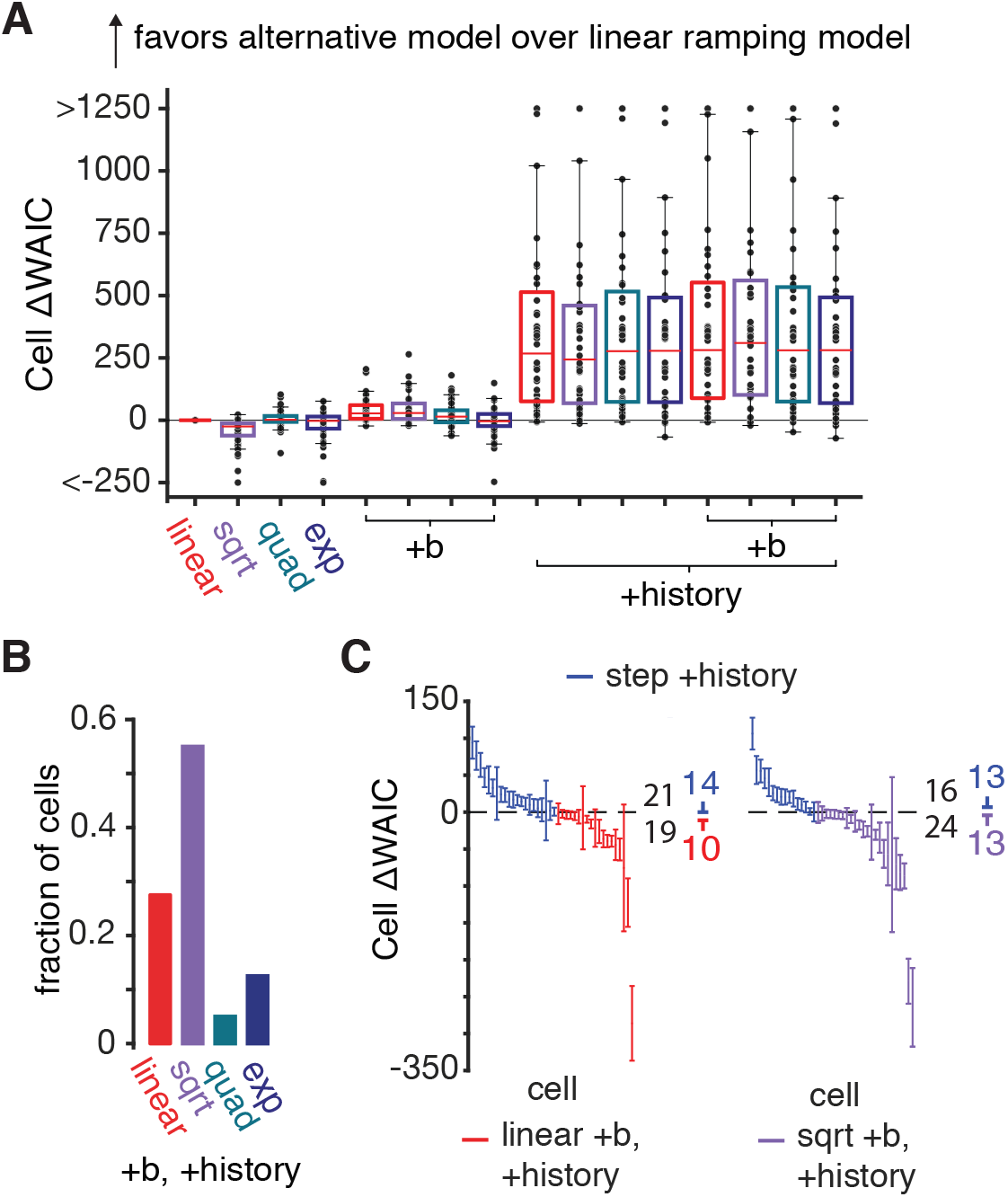
Comparison of extended ramping models. **A**. Comparison of nonlinear ramping models with non-zero baseline firing rates and spike-history against the classic (linear) ramping model. Positive values indicate improvements relative to the classic linear ramping model in terms of WAIC difference. The box color indicates the choice of nonlinearity and “+b” refers to models with a non-zero baseline. **B**. The fraction of cells for which each model (all with non-zero baseline and spike-history) achieved the best WAIC, showing that the square root nonlinearity performed best for more than half the population. **C**. WAIC differences between the stepping model with spike-history and the linear (*left*) and square root (*right*) ramping models with baseline and spike-history. See Fig. 8 for comparable analysis using cross-validation instead of WAIC.

As the linear and square root ramping models with non-zero baseline and spike-history were the best-performing among all models with continuous latents, we performed a direct comparison with the stepping model with spike-history. These extended ramping models closely matched the performance of the stepping model with spike-history (Figures 4C and 8C). The square root ramping model with non-zero baseline and spike-history achieved better WAIC than the stepping model with spike-history for more than half of the cells, although they each had an equal number of cells with WAIC differences more than the standard error from zero.

### 2.4 Inferred single-trial trajectories of nonlinear ramping models are more discrete

In the previous section, we showed that the linear and square root ramping models with non-zero baseline outperformed the linear ramping model for the majority of cells. We therefore investigated how these extensions affected different aspects of the model behavior. First, we observed that including a non-zero baseline helped to capture the steep initial decrease in the average firing rate for negative-coherence trials via steeper negative drift rates, which was missing for the simpler ramping model (Figure 5A-B and Figure 1C).

**Figure 5.**
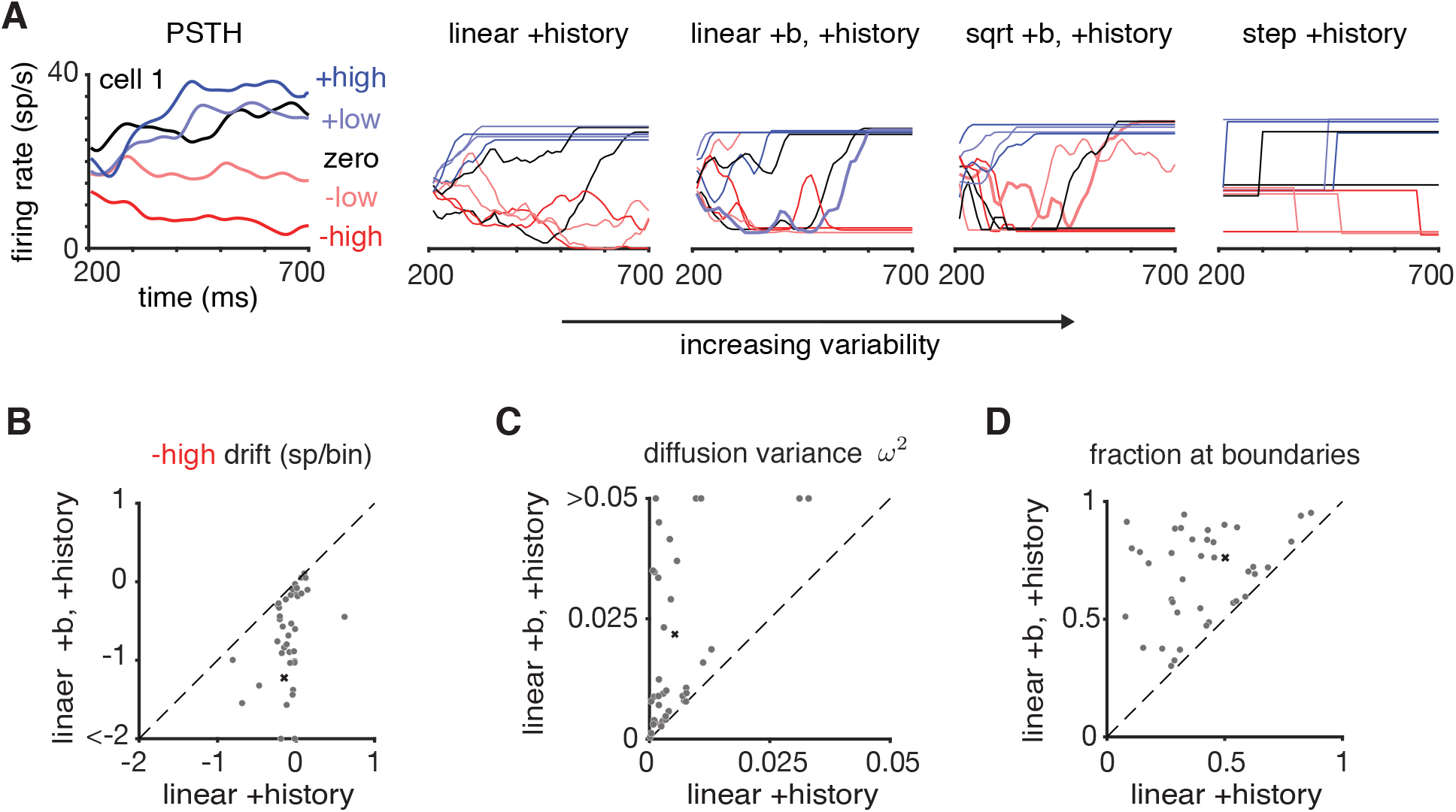
Analysis of latent firing rates under different latent dynamical models. **A**. Coherence-conditional PSTHs for example cell #1 (left) and simulated latent firing rate paths from fitted nonlinear ramping models and stepping model (right). The data were simulated using samples from the posterior. The linear and square root ramping models with non-zero baseline and spike-history can produce latent trajectories with high variability. The bolded trajectories are simulated trajectories that quickly evolved from the baseline to the upper boundary. **B-D**. Scatter plots comparing latent firing rate dynamics under the ramping model with and without a non-zero baseline. Each point corresponds to a pair of model fits for a single neuron. **B**. Drift rates for the highest negative-coherence stimulus are more negative under a ramping model with non-zero baseline than without. This indicates that adding a non-zero baseline allows the firing rate to ramp downwards more rapidly for negative-coherence motion (Zylberberg and Shadlen, 2016). **C**. Diffusion variance increased with the addition of the non-zero baseline. **D**. Fraction of time the simulated latent firing rates were equal to the baseline rate or upper absorbing boundary. See Figure 9 for comparable analyses with the square root ramping model with non-zero baseline.

Second, we observed that single-trial trajectories sampled from the models with non-zero baseline looked more discrete, exhibiting more rapid jumps to maximal or minimal firing rate, than trajectories from the linear ramping model without baseline (Figure 5A). Adding a non-zero baseline led to an increase in diffusion variability for most of the cells (Figure 5C), which allowed for larger changes in firing rate between time bins. Simulated trajectories also spent larger fractions of the time at the lower and upper boundaries compared to the models with a zero baseline (Figure 5D). Because the lower bound is non-absorbing, the firing rate can hit the lower bound and still evolve to the upper bound during the course of the trial; examples of these trials are shown in bold in Figure 5A. The effects of increased variability and increased time at minimal or maximal firing rates were also observed in the model with square-root nonlinearity and non-zero baseline when compared to the linear model (Figure 9).

Overall, these findings suggest that including a non-zero baseline rate can improve the ramping model in multiple ways. It does help the ramping model capture strong negative going rates (Zylberberg and Shadlen, 2016). However, with these modifications, the ramping model produced highly variable latent trajectories, which transitioned rapidly to minimal or maximal rates, making them qualitatively less similar to the gradually drifting rates expected from a perfect accumulator. Put simply, inclusion of a non-zero baseline moved both linear and nonlinear ramping models closer to discrete dynamics.

### 2.5 Generalization to two additional decision-making tasks

To see if the results for this dataset generalized to recordings from LIP in other direction discrimination tasks, we repeated a subset of our analyses on recordings during a discrete-pulse accumulation task (Yates et al., 2017) and a reaction-time (RT) task (Roitman and Shadlen, 2002; Section 4). The trial-averaged responses of neurons in both tasks resembled gradual ramps (Figure 6). However, statistical comparison of stepping and ramping models yielded results consistent with our findings above (Figures 6 and 10). For both additional datasets, spike-history filters improved the fits of both ramping and stepping models. The linear ramping model performed better with inclusion of a non-zero baseline, and the square-root ramping with non-zero baseline model outperformed the linear ramping with non-zero baseline model (both with history) for the majority of units in both datasets: 65/115 in the discrete-pulse task and 9/16 in the RT task. The stepping model with spike-history outperformed the linear ramping and square root ramping models with non-zero baseline and spike-history for a large majority of units in the discrete-pulse task. In the RT task, the models were evenly split, with models in the ramping class favored for cells with WAIC differences greater than the standard error from zero.

**Figure 6.**
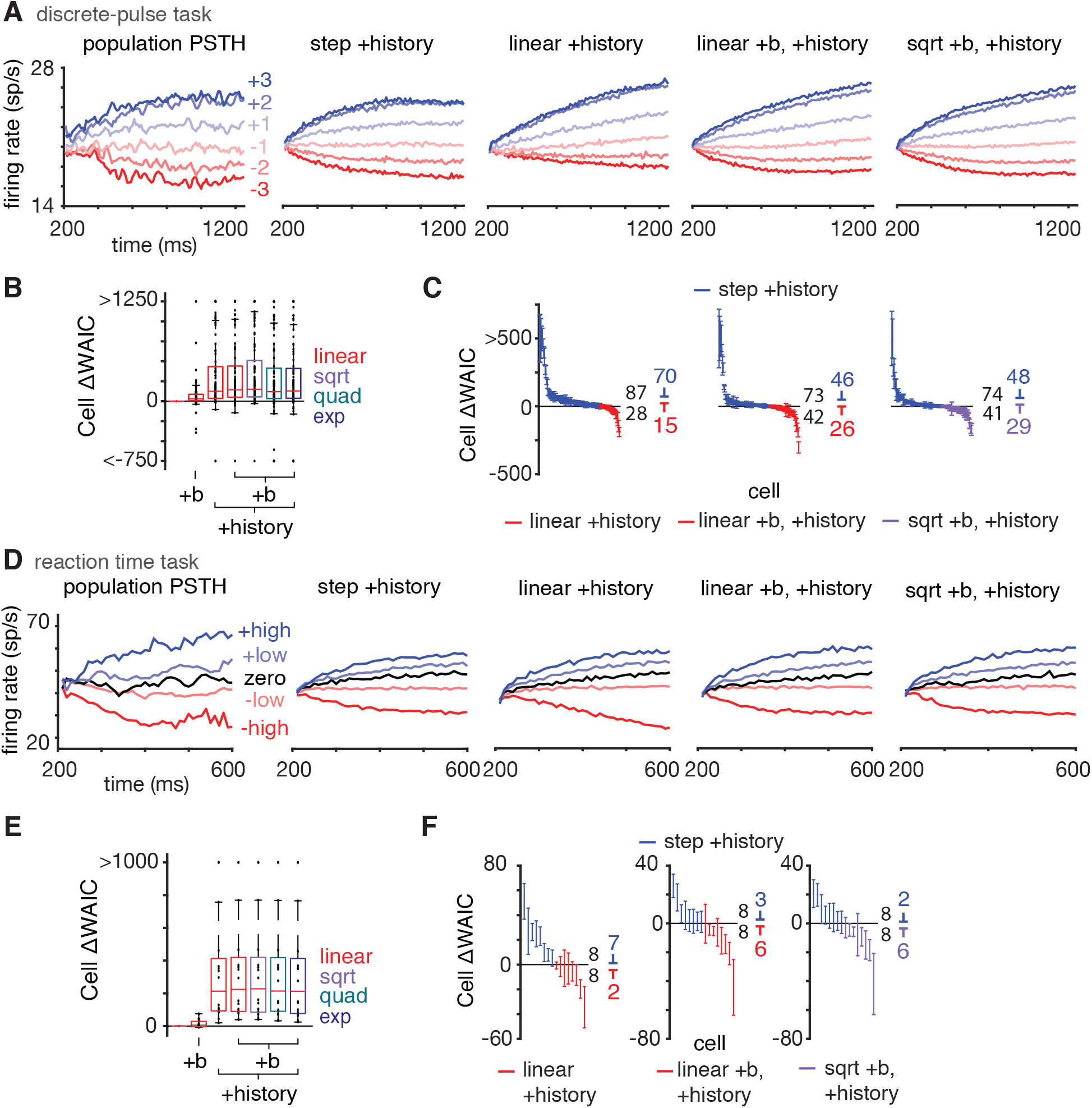
Analyses of neural data from two additional decision-making tasks. **A**. Population PSTHs from LIP neurons and simulated data from fitted models in a discrete-pulse task (Yates et al., 2017). **B**. Model comparison of extended nonlinear ramping models and classic linear ramping model using WAIC. **C**. The stepping model with spike-history achieved higher WAIC differences for the majority of cells. **D**. The population PSTH and simulated PSTHs from the fitted models, aligned to motion onset, in a reaction-time (RT) task (Roitman and Shadlen, 2002). **E**. Spike-history and non-zero baseline firing rates improved ramping model performance, but different nonlinearities performed equally well. **F**. Comparison of the stepping model and different ramping models, all with spike-history. See Figure 10 for comparable analyses using cross-validation.

It is worth summarizing the comparison between the stepping model and the square root ramping model with non-zero baseline (both with spike-history), as neither model outperformed the other across all datasets. Between the two models, the stepping model with spike-history was favored for slightly less than half of cells in the variable duration task (16/40), a majority of cells in the discrete-pulse task (73/115), and exactly half of cells in the RT task (8/16). However, for cells with WAIC differences greater than the standard error from zero, the stepping model with spike-history was favored for exactly half of cells in the variable duration task (13/26), a majority in the discrete-pulse task (46/72), and a minority in the RT task (2/8). In this view, across datasets these two models achieved relatively equal performance.

## 3 Discussion

Our study strengthens the evidence for discrete-state models of the single-trial dynamics of many LIP cells during decision-making. Importantly, we found that LIP dynamics are heterogeneous, with discrete stepping and continuous diffusion-to-bound models both accounting for a substantial fraction of neurons (Meister et al., 2013; Park et al., 2014; Latimer et al., 2015). Our findings are supported by dynamical models that account for spiking autocorrelation and that allow for nonlinear mappings from diffusion-to-bound to firing rate in the ramping model. We obtained the same results when using a fully Bayesian information criterion and leave-one-out cross validation.

Although we have significantly extended the models considered in Latimer et al. (2015), which allowed for more accurate descriptions of spike trains in LIP, there are a variety of other proposed extensions that we have not yet explored. For example, the ramping model could use a random start time to the diffusion process on each trial (Churchland and Kiani, 2016). We speculate that the ramping model has more flexibility to handle small deviations in the onset of a diffusion-to-bound process because it has continuous latent states that place mass over all possible trajectories in the latent space, although we have not performed this comparison explicitly. Next, one could formulate a ramping model in which negative drifting rates are stopped by a competing accumulator on each trial, rather than a non-zero baseline that is constant across trials (Mazurek et al., 2003; Zylberberg and Shadlen, 2016; Latimer et al., 2017). However, this would require a two-dimensional latent diffusion process, which would be computationally more demanding than the one-dimensional models we have considered here, and might prove more difficult to identify with single-neuron data. Fitting such a model using multi-neuron recordings therefore presents one promising direction for future work. Finally, one could extend the stepping model to a general hidden Markov model, allowing for more than three discrete firing rates, with more flexible transition dynamics that allow for more than one transition per trial (Bollimunta et al., 2012). Such a model would have more flexibility than the stepping model we considered, which might allow better generalization to alternate tasks (Janssen and Shadlen, 2005; Yang and Shadlen, 2007; Kira et al., 2015; Morcos and Harvey, 2016).

Our findings appear to contradict a recent study from Zhao and Kording (2018), which reported that the best model of LIP responses, according to a cross-validation analysis, was a model with a constant firing rate on every trial. Although the specific models differed from those we have considered here in multiple ways, we believe the discrepancy is likely due to the fact that Zhao and Kording (2018) treated latent firing rates as parameters to be estimated, instead of marginalized or integrated over. This resulted in models with one fitted parameter per trial (a step time, ramp slope, or constant firing rate), making for hundreds of parameters per neuron, which is far more than the models we have considered here. We suspect that this approach therefore suffered from overfitting, leading to the dubious conclusion that firing rates are constant over time (a result that is inconsistent with the basic ramping apparent in trial-averaged activity). We have shown here that WAIC and cross-validation give virtually identical results when integrating over the unobserved latent firing rates. Nonetheless, the suggestion from Zhao and Kording (2018) that baseline firing rates may vary stochastically across trials is interesting, and consistent with recent findings about spike count variability (Goris et al., 2014; Charles et al., 2018). Incorporating slow changes in gain or excitability over trials therefore represents an additional promising direction for future work.

Some authors have raised the concern that animals performing the task are adopting a strategy that involves integration over a shorter time period than the entire trial, which could produce discrete-looking neural dynamics even in neurons that are accumulating evidence (Shadlen et al., 2016). Further, alternative strategies without accumulation can also match some behavioral features of evidence accumulation (Ditterich, 2006b; Stine et al., 2018). We recognize the ambiguity in determining behavioral strategy, and in future work we expect that including the time-varying evidence stream in the models could help identify the behavioral strategies used by the animals.

The GLM framework we have used to incorporate spike-history effects could naturally be expanded to include regressors for experimental variables related to the stimulus or behavior of the animal. A worthwhile future direction would be to fit latent variable models with GLM regressors in tasks with structured stimuli, to better disentangle latent dynamics from sensorimotor variables that affect neural activity on single trials (Brunton et al., 2013; Hanks et al., 2015; Morcos and Harvey, 2016; Katz et al., 2016; Scott et al., 2017; Yates et al., 2017; Huk et al., 2017). As behavioral paradigms become richer, and as the numbers of recorded neurons increase, we expect that population latent variable models with regressors and GLM outputs will provide a powerful framework for studying the neural computations underlying sensory decision-making in a wide variety of tasks and brain areas.

## 4 Methods

### 4.1 Data

We analyzed the responses of LIP cells during three motion-discrimination tasks. Our primary analyses were performed on the responses of 40 LIP cells (single-units) recorded from two rhesus monkeys during a variable-duration random dot motion task, originally described in Meister et al. (2013). In the task, a random dot motion stimulus was presented for durations uniformly drawn in the range 500 to 1000 ms after a variable delay. The dot motion coherence, or the expected percentage of dots moving in the true direction at each time point, on each trial was taken from the set of values: 0.0, 3.2, 6.4, 12.8, 25.6, or 51.2%. The monkey reported its estimate of the dot motion direction via a saccade to one of two targets. One saccade target was placed in the response field of the neuron (“in-RF”) while the other was placed outside of the response field (“out-RF”). The animal had to wait for 500 ms after the stimulus was extinguished before it could indicate its choice. The original study recorded from 80 LIP neurons and the 40 LIP cells used in this study were the 40 most choice-selective responses during the period 200–700 ms after motion onset, determined by the *d^′^* criterion (Latimer et al., 2015). For analysis, the trials were grouped into five coherence levels: zero included 0% trials, positive/negative low included 3.2%, 6.4%, and 12.8% trials, and positive/negative high included 25.6% and 51.2% trials, where positive motion is towards the target in the response field. The coherence-dependent PSTHs for each unit in the variable duration task were smoothed using a Gaussian filter with 20 ms standard deviation.

We analyzed two additional datasets. The first consisted of 16 LIP cells (single-units) recorded from two rhesus monkeys during a reaction-time (RT) version of the random dot motion task (Roitman and Shadlen, 2002), where the monkey chooses when to respond. The details for the selection of the 16 LIP cells are in (Latimer et al., 2015). We also divided the trials in this dataset into the five levels described above. We included spikes starting at 200 ms after stimulus onset and up to 50 ms before the saccade for analysis. The final spike bin contained the time point 50 ms before the saccade and we included all spikes that fell into this bin. We only included trials in which we had 100 ms of data in this period (Latimer et al., 2015).

The second additional dataset consisted of 115 LIP units (single-and multi-units) from two rhesus monkeys performing a discrete-pulse accumulation task (Yates et al., 2017). In this task, the animal viewed a set of Gabor patches that either flickered or drifted during seven discrete portions of the trial (pulses). In each pulse, all of the drifting Gabor patches moved in the same direction. The task of the animal was to report the net motion direction across the seven pulses with a saccade to one of two targets. In this task, the net motion levels did not map directly to discrete-coherence levels, as each trial could have a different amount of net pulses in either direction. Therefore, we partitioned the data for each experimental session into six levels by sectioning the net pulses in each direction into thirds. We selected neurons with *d^′^* statistic magnitudes larger than 0.2 for analysis (Yates et al., 2017).

### 4.2 Models

#### 4.2.1 Ramping model with history

In the ramping model, the firing rate is linked to a latent diffusion process (Latimer et al., 2015). In the following, we describe the generative model. Individual trajectories in the latent space are initiated with a sample from a Gaussian with mean *x*_0_ and variance *ω*^2^. The trajectory then evolves according to drift-diffusion dynamics with a diffusion variance *ω*^2^ and drift *β_c_*. The drift *β_c_* depends on the coherence *c* of the current trial. The latent trajectory is scaled by a factor of *γ* and passed through a nonlinearity *f*(*x*) to map it to a positive firing rate space. In the linear ramping model, the nonlinearity is the softplus function *f*(*x*) = log(1 + exp(*x*)). The output of the nonlinearity is multiplied by history dependence *g_t_* such that the firing rate in spikes per second is *λ_t_* = *f*(*x_tγ_*) *g_t_*. If the trajectory crosses an absorbing upper boundary at 1 in the latent space then the firing rate is held fixed for the remainder of the trial at the boundary rate. The generative model for a trial of length *T* is 
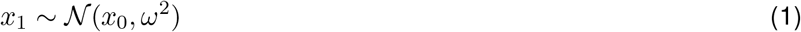

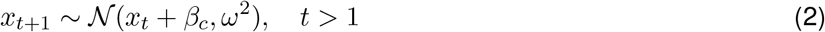

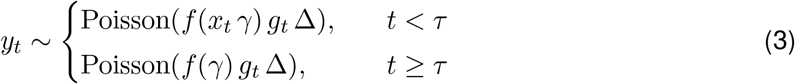
where *τ* is the first time bin that *x_t_* ≥ 1 (otherwise *τ* = *∞*) and Δ = 0.01*s* is the bin size. The bin size is equal to one frame of the stimulus in Meister et al. (2013).

The history dependence modulates the firing rate through a multiplicative interaction. At time *t*, the history dependence *g_t_* is the exponential of the weighted sum of the previous *H* bins of spiking activity of the neuron 
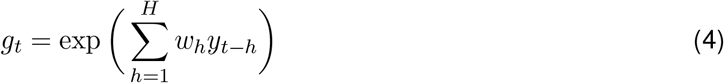
with w = [*w*_1_, · · ·, *w_H_*]^T^ a vector of weights. In the models without history *g_t_* = 1 for all *t*. We used *H* = 10 bins for 100 ms of history dependence.

The ramping model parameters are Θ = {*β*_1_:_*C*_, *x*_0_, *ω*^2^, *γ*, w} where *C* is the number of coherence levels. The latent variables in the ramping model x are the latent diffusion trajectories for each trial.

#### 4.2.2 Nonlinear ramping models

The nonlinear ramping models are linked to the latent diffusion process of the ramping model through alternative nonlinearities. We used three alternative nonlinearities: a soft square root *f*(*x*) = log(1 + exp(*x*))^1/2^ (sqrt), a soft quadratic *f*(*x*) = log(1 + exp(*x*))^2^ (quad), and an exponential *f*(*x*) = exp(*x*) (exp). The parameters and latent variables of the nonlinear ramping models are unchanged from the ramping model.

#### 4.2.3 Non-zero baseline firing rate

In the linear and nonlinear ramping models with non-zero baseline rates, the output of the nonlinearity is shifted by a positive baseline firing rate parameter *b* before multiplication with the history term (Zylberberg and Shadlen, 2016; Latimer et al., 2017) such that the firing rate is *λ_t_* = (*f*(*x_tγ_*) + *b*) *g_t_*. The generative model is 
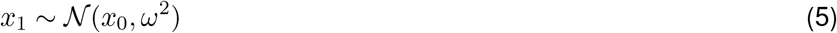

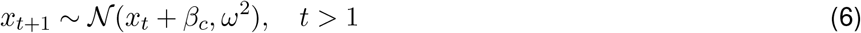

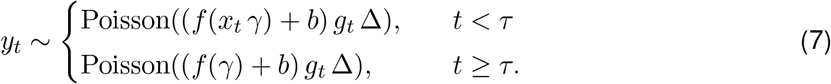

The parameters of the ramping and nonlinear ramping models with non-zero baseline are Θ = {*β*_1:*C*_, *x*_0_, *ω*^2^, *γ, b*, w}, where *C* is the number of coherence levels. The latent variables are x.

#### 4.2.4 Stepping model with history

In the stepping model, the initial firing rate starts at a state *α*_0_. During the trial, the state can either remain constant or it can switch to one of two other states, a down state *α*_1_ or an up state *α*_2_ (Latimer et al., 2015). The step direction *d* is sampled from a Bernoulli distribution such that the probability of a step to *α*_2_ is *ϕ_c_* and the probability of a step to *α*_1_ is 1 − *ϕ_c_*. The step time *z* is drawn from a negative binomial (NB) distribution with a shape parameter *r* and coherence-dependent mean step time *m_c_*. Both the step direction and step time vary from trial to trial. The stepping model firing rate is the product of the state and the spike-history dependent gain *g_t_*. The generative model for a trial of coherence *c* is 
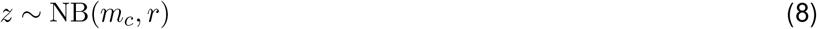

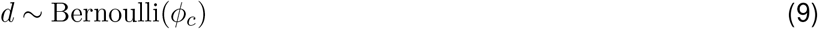

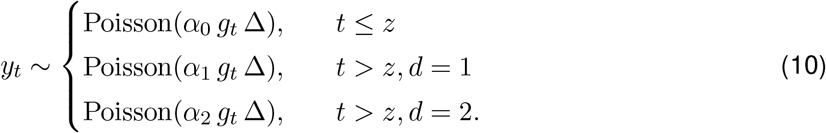

The stepping model parameters are Θ = {*α*_0_, *α*_1_, *α*_2_, *m*_1:*C*_, *ϕ*_1:*C*_, *r*, w}. The bin size is Δ = 0.01*s*. The history-dependence has the same parameterization as the ramping models. The latent variables in the stepping model are the step times and step directions on each trial. This stepping model is a reparameterization of the stepping model in (Latimer et al., 2015), which used scale parameters *p_c_* instead of *m_c_*, where 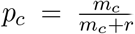. We used the mean step time parameterization because the parameters *r* and *m_c_* are less correlated than *r* and *p_c_*, which improved mixing in the MCMC algorithm described below.

### 4.3 Model Inference

#### 4.3.1 Prior Distributions

We used the following priors on the parameters of the ramping and nonlinear ramping models 
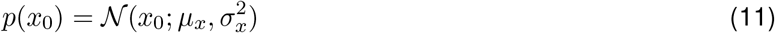

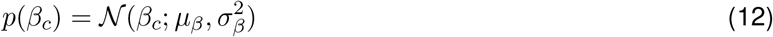

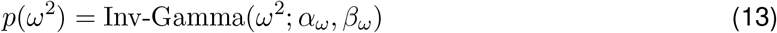

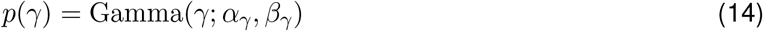

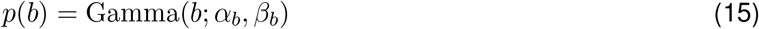

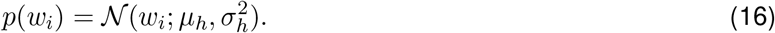

For all models, the priors on the diffusion drifts and variance were *µ_β_* = 0, *σ_β_*= 0.1, *α_ω_* = 1.1, and *β_ω_* = 1e−3. In models with a zero baseline, we set *µ_x_* = 0 and σ*_x_* = 10. In models with a non-zero baseline, we set *µ_x_* = 0.5 and σ*_x_* = 0.5 and used *α_b_* = 1 and*_b_* = 0.01 for the prior on the baseline parameter. The prior on the bound height varied for each nonlinearity. We used *α_γ_* = 2 and *β_γ_* = 0.05 for the softplus, *α_γ_* = 1 and *β_γ_* = 1e−4 for the soft square root, *α_γ_* = 3 and *β_γ_* = 0.5 for the soft quadratic, and *α_γ_* = 3 and *β_γ_* = 3 for the exponential. The parameters for the prior on the history weights were *µ_h_* = 0 and 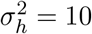.

The priors on the stepping model with history were 
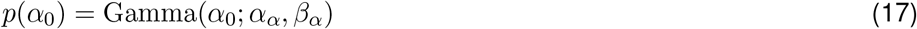

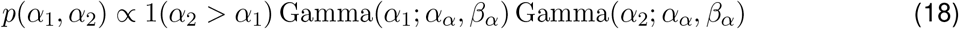

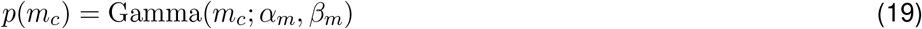

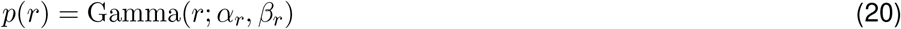

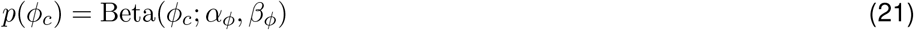

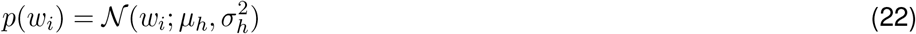
where 1(·) is the indicator function. The joint prior on the rates *p*(*α*_0_, *α*_1_, *α*_2_) = *p*(*α*_0_)*p*(*α*_1_, *α*_2_) enforces identifiability and we set *α_;_* = 1 and *β_α_* = 0.01. The prior over the mean step times was*_m_* = 2 and *β_m_* = 0.02. This prior has a peak at 50, a mean of 100, and it places significant mass over a broad range of *m*. The prior over the step direction probabilities was uniform over [0,1] with *α_ϕ_* = 1 and _β*ϕ*_ = 1. The prior over *r* used *α_r_* = 2 and *β_r_* = 1. The history weights had the same prior as in the ramping model with *µ_h_* = 0 and 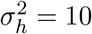. For the stepping model without history, the priors were those specified in (Latimer et al., 2015).

#### 4.3.2 MCMC Overview

We used Markov chain Monte Carlo (MCMC) methods to obtain approximate samples from the posterior of the model parameters Θ given the data y, *p*(Θ|y). Specifically, we used MCMC to approximately sample from the joint posterior of the parameters and the latents *p*(Θ, x|y) and ignored the samples of the latents to obtain samples from *p*(Θ|y).

The following is an overview of the MCMC methods we used to obtain samples from *p*(Θ|y) (Latimer et al., 2015). First, we sampled a value for each latent variable given the parameters and the observed spike counts from the distribution *p*(x|Θ, y). Then, conditioned on the new value of the latent variables and the data, we sampled new parameter values from *p*(Θ|x,y). By repeating this procedure many times we obtained samples from the distribution *p*(Θ, x|y). We marginalized over x to obtain the posterior distribution over the parameters *p*(Θ|y) by simply discarding the values of x. For each MCMC simulation, we simulated a chain of 60000 samples from *p*(Θ|y). We discarded the first 10000 samples from the chain as a burn in period. We kept every fifth sample afterwards, leaving *S* = 10000 samples from the posterior distribution over the parameters. For each posterior sample {Θ^*s*^}_s= 1:*S*_, we computed the likelihood of the data given the posterior sample *p*(y|Θ^*s*^) by marginalizing over the latent variables. In the ramping models, we used 5000 Monte Carlo samples of the latent trajectories given Θ^*s*^ to compute the likelihood. In the stepping model, we performed the marginalization by integrating the step times and step directions on a grid (Latimer et al., 2015).

For the variable duration dataset, we simulated two MCMC chains for the ramping, linear ramping with non-zero baseline, square root ramping with non-zero baseline, and stepping models (all with spike-history) to examine convergence in the MCMC chains before comparing these models. We assessed convergence using the potential scale reduction factor (PSRF) convergence diagnostic (Gelman et al., 2013) on the trial likelihoods from the two chains. We chose to monitor the convergence of the likelihoods because our model comparison is based solely on the likelihoods. If the diagnostic indicated that the two chains had not converged to the same likelihood distribution (PSRF > 1.1), we simulated additional chains until we obtained two chains that passed the diagnostic. This required increasing the number of burn in samples for a few cells.

#### 4.3.3 MCMC for Ramping and Nonlinear Ramping Models

The MCMC sampling procedure for the ramping and nonlinear ramping models proceeded as follows. We first initialized the parameters to Θ^(1)^. We set β_1:*C*_ by sampling *C* values from the distribution 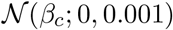 and sorting the values in the order of the coherence levels. We set the initial bound height *γ* to be a sample from a Gaussian distribution with mean equal to the average spike rate in the final time bin of in-RF choice trials and with unit variance. We sampled the initial *x*_0_ from a Gaussian with mean equal to the average spike rate in the first time bin divided by the initial *γ* and with standard deviation 0.01. We constrained the initial *x*_0_ to be in [0.1, 0.9]. We sampled the initial variance *ω*^2^ uniformly in the range [5e−4, 5e−3]. We sampled the initial history weights 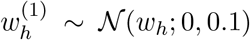. With a non-zero baseline, we sampled the initial baseline parameter 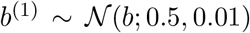 and also subtracted the baseline from the mean of the distribution for sampling *γ*.

After initializing the parameters, we alternated between sampling the latent diffusion paths conditioned on the current parameters and sampling new values of the parameters conditioned on the previous latent path. Formally, we obtained the *s*^th^ sample, for *s >* 1, with 
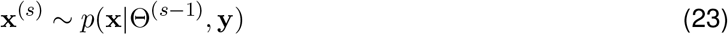

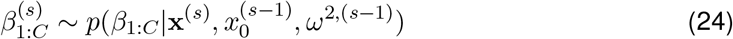

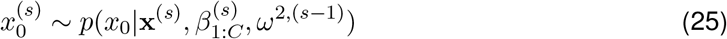

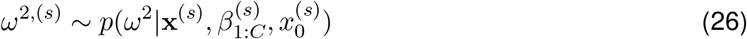

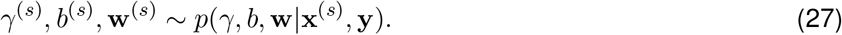

For step (23), we used a particle filter to estimate the distribution of latent paths below the boundary and the distribution of boundary crossing times (Latimer et al., 2015). Given those two distributions, we used a backwards sampling scheme to sample the latent paths x^(*s*)^. We modified the firing rate observation likelihood in this step for each model to include the appropriate nonlinearity, baseline, and history dependence. We exploited conjugacy in steps (24), (25), and (26) for Gibbs steps, which were identical to those presented in Latimer et al. (2015).

For the final step (27), we used a manifold Metropolis-adjusted Langevin (MMALA) step to jointly sample the parameters *θ* = [ *γ, b*, w]^T^ (Girolami and Calderhead, 2011; Latimer et al., 2015). The vector of parameters has dimension *J* = 2 + *H*, where *H* is the number of history weights. In the following derivation, for models with a subset of the parameters *θ*, the terms unrelated to the subset of parameters are disregarded. Each Metropolis step consisted of sampling a new value of the parameters *θ*^*^ from a proposal distribution *q*(*θ*^*^|(*θ*^(*s*−1)^, y, x^(*s*)^) and accepting the newly sampled values (that is, set *θ*^(*s*)^ = *θ*^*^) with probability 
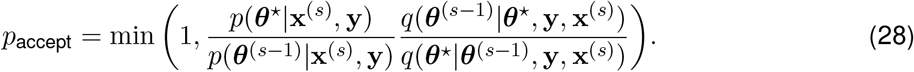

If the proposed values were not accepted then we set *θ*^(*s*)^ = *θ*^(*s*−1)^. The proposal distribution used the gradient of the log likelihood plus log prior ∇_*θ*_*ℒ*(*θ*) and the Fisher information matrix plus Hessian of the log prior G(*θ*) 
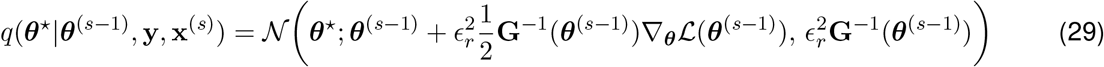
where 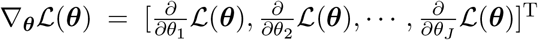. The step size *∊_r_* was initialized to 0.05 and gradually increased to 1 during the burn in period.

We define the firing rate function 
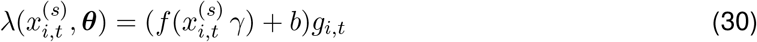
that is in general a function of *γ*, the baseline parameter *b*, the sampled latent path, and the history for trial *i* and time *t*. The log likelihood plus log prior is a sum over *N* trials and *T_i_* time points on each trial 
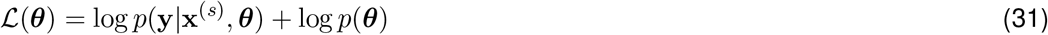

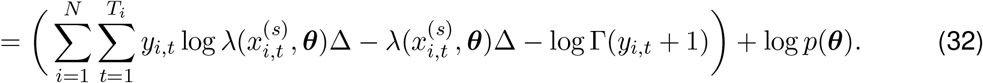

The gradient of *ℒ*(*θ*) is defined by the derivatives with respect to each parameter*θ_j_* 
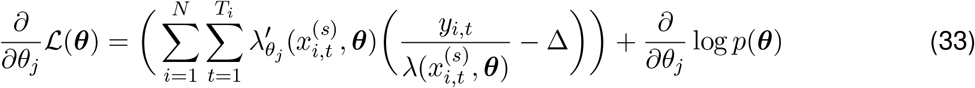
where 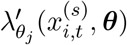 is the derivative of the rate function with respect to *θ_j_*. The *J* × *J* matrix G(*θ*) is 
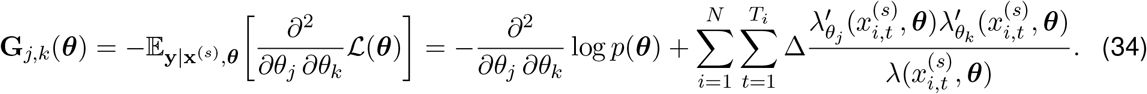

The derivatives 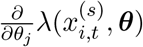 for each parameter are 
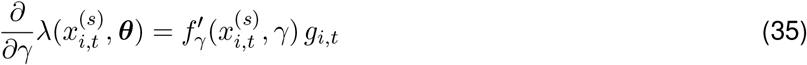

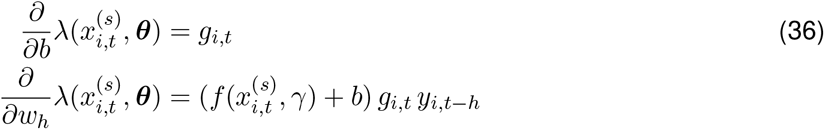
where 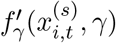 is the derivative of each nonlinearity with input 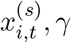, *γ* with respect to *γ* 
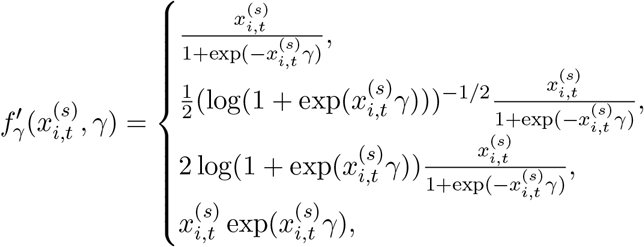

The derivative and second derivative of the log prior on *γ* are 
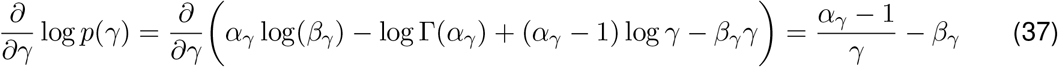

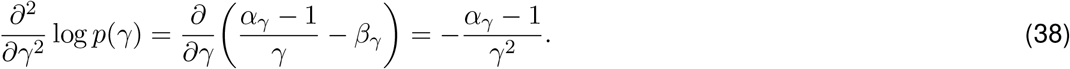

Similarly, for the baseline *b* these quantities are 
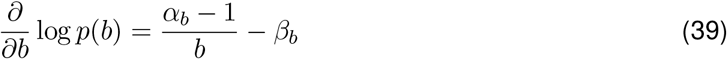

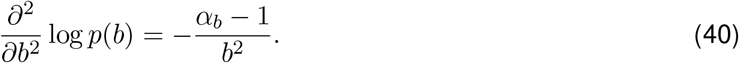

The first two derivatives of the log likelihood of the history weights *w_h_* are 
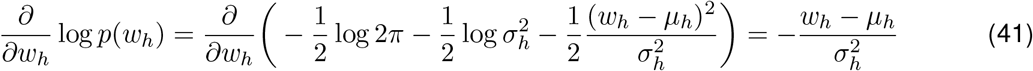

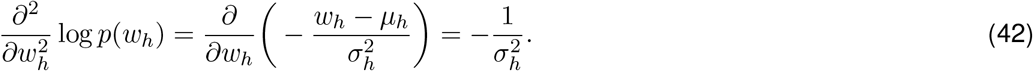

Each of the priors are independent and therefore the prior terms contributing to the off-diagonal elements of G_*j,k*_(*θ*) are zero.

#### 4.3.4 MCMC for Stepping Models

In the stepping model we alternated between sampling the latent step times z and directions d and sampling the parameters of the model Θ. We first initialized the parameters to Θ^(1)^. We set 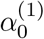 to the firing rate in the first time bin, 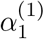 to the firing rate in the final time bin of out-RF choice trials, and 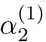 to the firing rate in the final time bin of in-RF choice trials. We then added Gaussian noise to each 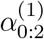. We sampled 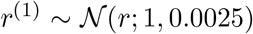, 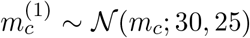, 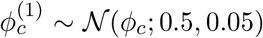, and 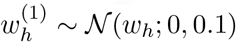.

After initialization, we performed the following sequence of steps to obtain the *s*^th^ > 1 sample 
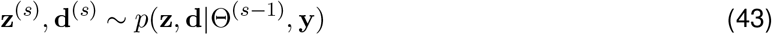

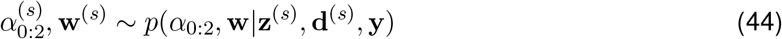

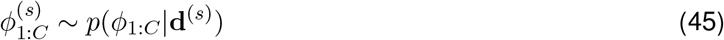

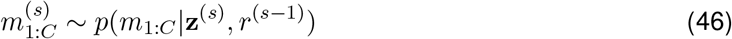

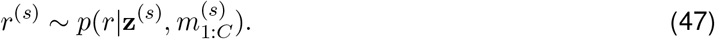

We sampled the step directions and step times (step 43) by sampling from the distribution computed on a grid, truncated at 1500 time bins (Latimer et al., 2015), with the history dependence included in the observation likelihood. We employed Beta-Bernoulli conjugacy to directly sample the step direction probabilities (step 45) using a Gibbs’ step.

##### Sampling the rates and spike history filters

We used an MMALA step in the stepping model to sample the rates *α*_0:2_ and history weights w with proposal distribution 
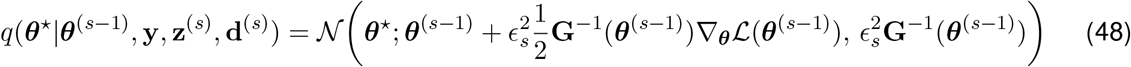
where *θ* = [*α*_0:2_, w]^T^. The step size *∊_s_* was gradually increased from 0.05 to 1 during the burn in period. The firing rate for trial *i* and time *t* is 
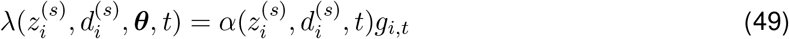
where 
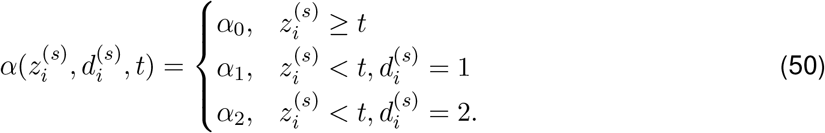

The MMALA step uses *ℒ*(*θ*), 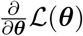, and G(*θ*). The log likelihood plus log prior is 
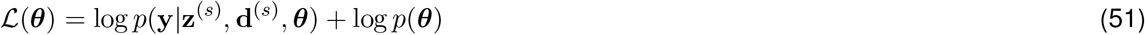

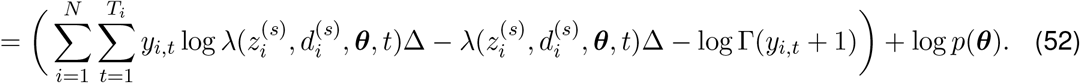

The gradient of the log likelihood plus log prior is 
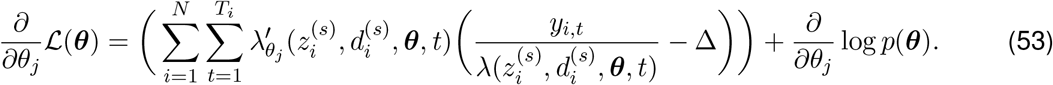

The elements of the Fisher information matrix plus the Hessian of the log prior are 
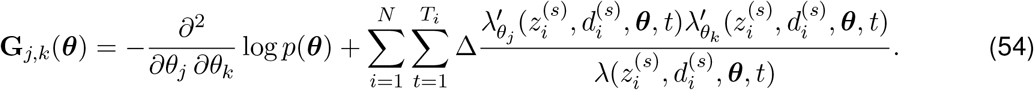

The derivative of the rate function given 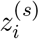 and 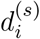 is 
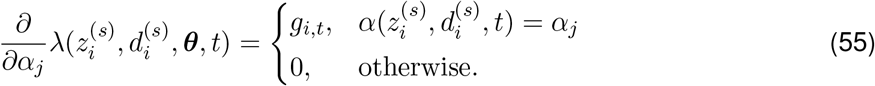

The derivative with respect to the history weights is 
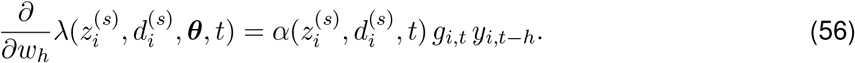

When evaluating the first and second derivatives of the log prior for the proposal distribution in (53) and (54), we used independent priors on ;_1_ and ;_2_ 
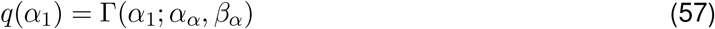

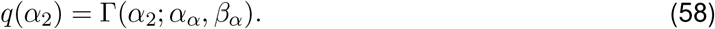

This simplifies computation of the gradient and Hessian for the proposal distribution. The derivatives of the log prior for *α*_0:2_ and w have the same form as and w in the ramping model.

We note that if across all trials the cell was never in state *α_j_* then 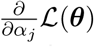 is zero and the row and column of G(*θ*) corresponding to *α_j_* is zero. This matrix must be nonsingular such that we can use its inverse in the proposal distribution. Therefore, if this occurred, although rare, we set the diagonal element of each zero row and column to one.

##### Sampling step time means and shape

We sampled the mean *m*_1:*C*_ and shape *r* parameters of the negative binomial distribution over the step times using Metropolis steps. The probability of a step time *z_i_* on trial *i* with coherence *c_i_* in terms of *m_ci_* and *r* is 
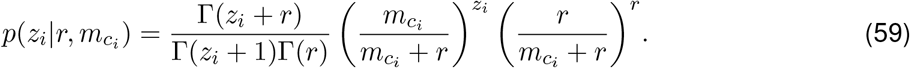

We alternated between sampling each *m_c_* conditioned on *r* and sampling *r* conditioned on each *m_c_*.

The proposal distribution for each *m_c_* is 
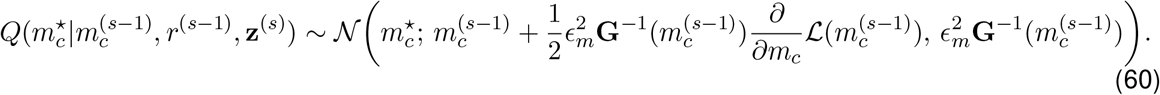

We gradually increased *∊_m_* from 0.05 to 1 during the burn in period. The log likelihood of *m_c_* plus log prior is the sum of the likelihoods of the step time *z_i_* for each trial with coherence *c* 
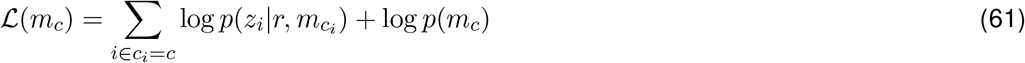

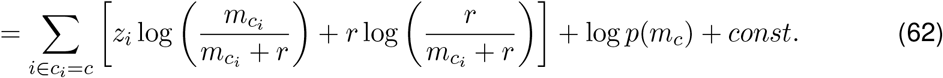

The derivative of the log likelihood of *m_c_* plus log prior with respect to *m_c_* is 
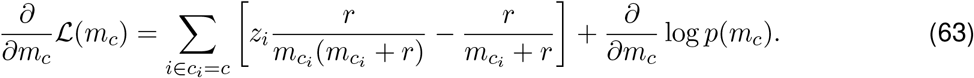

The Fisher information plus the Hessian of the log prior is 
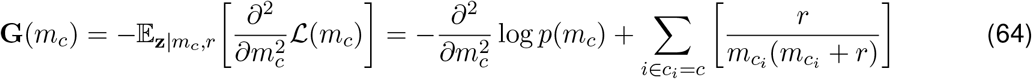
where we have used 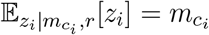.

The proposal distribution for *r* is 
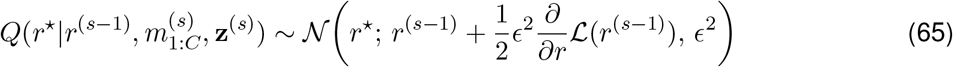
and the log likelihood of *r* plus log prior is a sum over all trials 
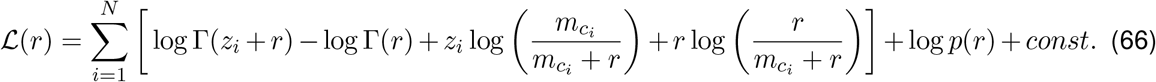

The derivative of the log likelihood plus log prior with respect to *r* is 
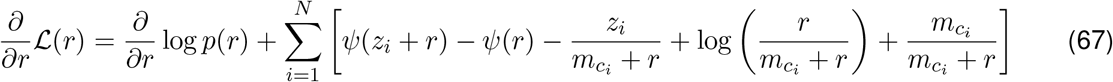
where *ψ* is the digamma function *ψ*(*r*) = ψ^′^(*r*)/ ψ(*r*). The step size *ϵ* was initialized to 0.075 and was adjusted throughout the burn in period, after which it was fixed.

### 4.4 Model Comparison: WAIC

We used the WAIC to compare the models (Watanabe, 2010; Gelman et al., 2014; Vehtari et al., 2017; Piironen and Vehtari, 2017). The WAIC estimates the expected generalization of a fit model to new data from the true data generating distribution. This corresponds to an estimate of how well the model would predict spike trains recorded on new trials. As generally the experimenter neither has access to unlimited data nor the true data generating distribution, the WAIC and other information criterion methods estimate the expected generalization using how well the model describes the in-sample data with a correction factor.

The WAIC is a function of the probability of the data given each posterior sample, {*p*(y_*i*_|Θ^*s*^)}*_s_*_= 1:*S*_, for each trial *i*. We used the formula in (Gelman et al., 2014) to compute the WAIC across *N* trials as 
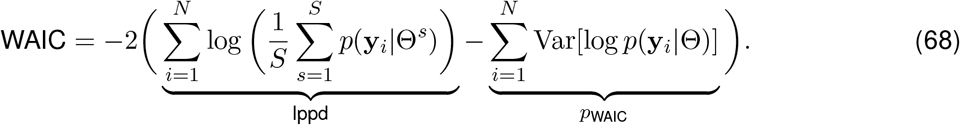

The first term is the log pointwise predictive density (lppd) and it describes how well the model predicts the data to which it was fit. A strength of the WAIC is the lppd averages over the posterior rather than conditioning on a point estimate of the parameters. The second term *p*_WAIC_ is a penalty that corrects the bias induced by estimating the expected generalization to new data from the lppd. The penalty term *p*_WAIC_ is computed for each trial as the variance of the log likelihoods of a trial across the posterior samples {Θ^*s*^}_*s*= 1:*S*_, and therefore is guaranteed to be non-negative because it is a sum of variances, another strength of the WAIC (Gelman et al., 2014). Additional advantages of the WAIC are theoretical results showing its asymptotic equivalence to Bayesian leave-one-out cross validation, its applicability to singular statistical models, and its computational efficiency when compared to leave-one-out cross validation (Watanabe, 2010; Gelman et al., 2014; Piironen and Vehtari, 2017). For all of these reasons we used it to compare the relative fits of the models.

In model comparison, we computed the WAIC difference between two models 
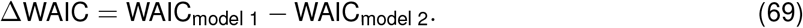

Since lower WAIC values are better, a positive difference favors model two while a negative difference favors model one. In some cases, we normalized the WAIC by the number of trials to put comparisons with differing numbers of trials on the same scale. We also considered the WAIC difference on subsets of trials by only summing across trials of certain conditions. We set the stepping model with spike-history as model 2 in model comparison with other models. Therefore, positive WAIC differences favor the stepping with spike-history model over the alternative model in these comparisons.

We quantified uncertainty in the model comparison using standard errors of the WAIC differences across trials 
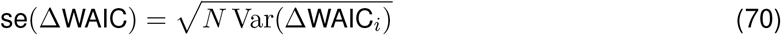
where ΔWAIC_*i*_ is the WAIC difference computed for trial *i*.

### 4.5 Simulated Data

We computed simulated latent trajectories and PSTHs from a fit model using the following procedure. For 40 random samples from the posterior over the parameters, we simulated a spike train for each trial conditioned on the pre-trial spiking activity and parameters of the model. For a few cells, simulated spike trains from the models with spike-history generated unrealistically large numbers of spikes due to self-excitation. We enforced realistic spike trains in these cases by setting the multiplicative history gain to unitary whenever the generated spike history effect was larger than the largest inferred history gain in the data. We averaged the simulated spike trains corresponding to each coherence to compute the coherence-dependent simulated PSTHs.

We computed the autocorrelation of the observed and simulated data for each neuron with the normalized autocorrelation function 
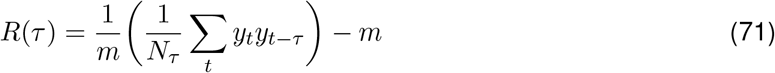
where *y_t_* is the spike count at time *t* and *m* is the mean spike count. The sum was computed over all valid time bins and *N_τ_* is the number of valid time bins.

For computing the fraction at the boundaries in (Figure 5), we used the following criterion. A time point *t* was classified as at a boundary if *x_t_* ≤ 0 or *x_t_* ≥ 1.

## Acknowledgements

This work was supported by the McKnight Foundation (J.W.P.), NSF CAREER Award IIS-1150186 (J.W.P.), and grants from the NIH (NEI grant EYE017366, to A.C.H., and NIMH grant MH099611, to A.C.H. & J.W.P.). David Zoltowski is supported by NIH grant T32MH065214. Jacob Yates is an Open Philanthropy Fellow of the Life Sciences Research Foundation. Kenneth Latimer is a Chicago Fellow.

## Author Contributions

Conceptualization, D.M.Z., K.W.L., A.C.H., and J.W.P.; Methodology, D.M.Z., K.W.L., A.C.H., and J.W.P.; Software, D.M.Z.; Formal Analysis, D.M.Z.; Investigation, J.L.Y.; Data Curation, J.L.Y.; Writing – Original Draft, D.M.Z. and J.W.P.; Writing – Review & Editing, D.M.Z., K.W.L., J.L.Y., A.C.H., and J.W.P.; Visualization, D.M.Z.; Supervision, J.W.P.; Funding Acquisition, A.C.H. and J.W.P..

## A Supplementary Figures

**Figure 7.**
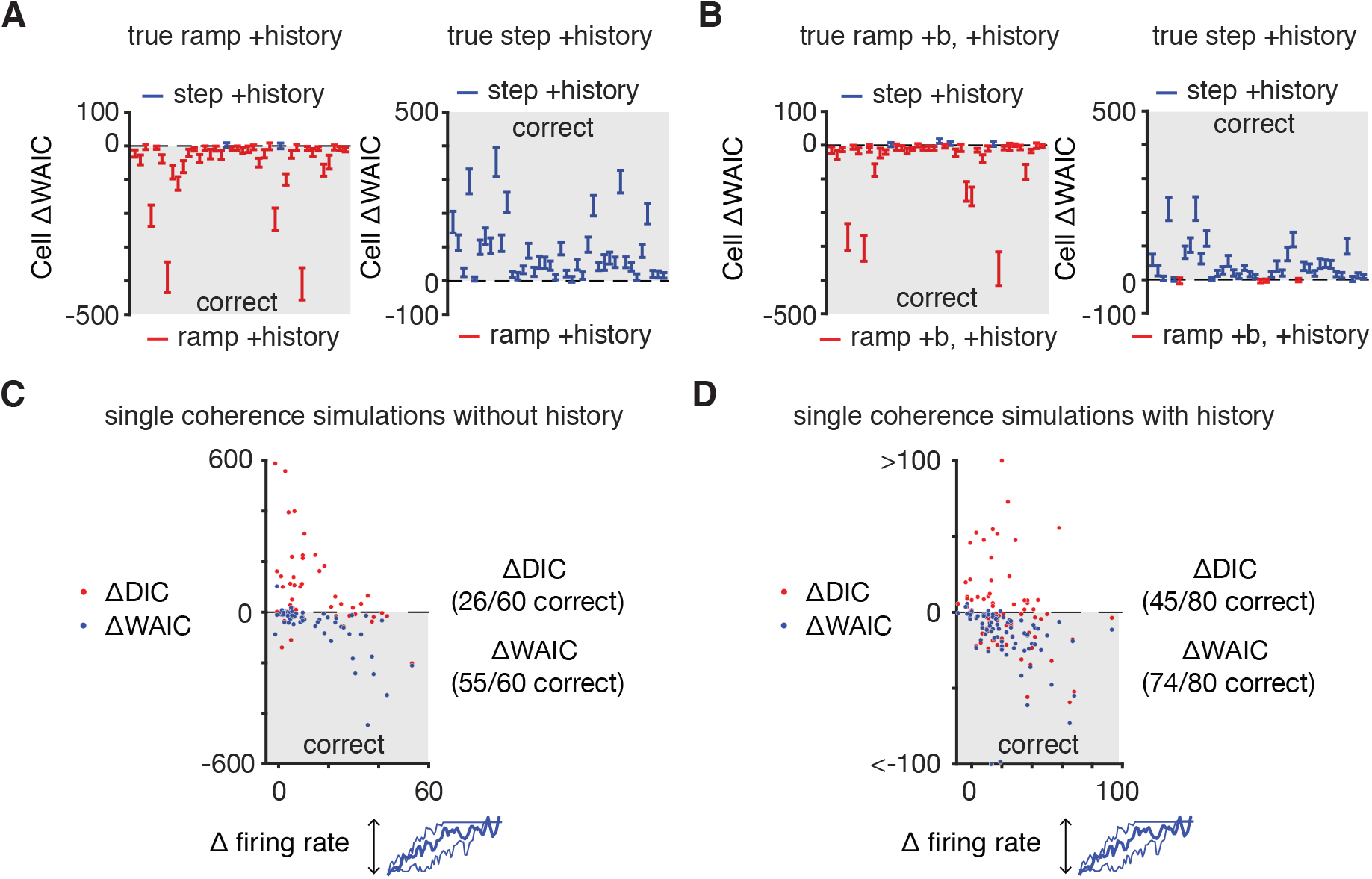
The WAIC correctly identifies simulated data from different models. Related to Figures 3 and 4. **A**. The WAIC reliably distinguished data from the ramping and stepping models with spike-history simulated from the mean parameters of the ramping (left) and stepping (right) models with spike-history for each cell. The number of simulated trials per motion coherence was matched to the true number for each cell. **B**. Same as **A**, except for simulated data from the ramping with non-zero baseline and history model. **C.-D**. Model comparison results from simulated single-coherence ramping data as a function of the change in firing rate from the beginning to the end of the trial using the DIC and WAIC. The WAIC correctly identifies data simulated from single-coherence ramping models (Chandrasekaran et al., 2016).

**Figure 8.**
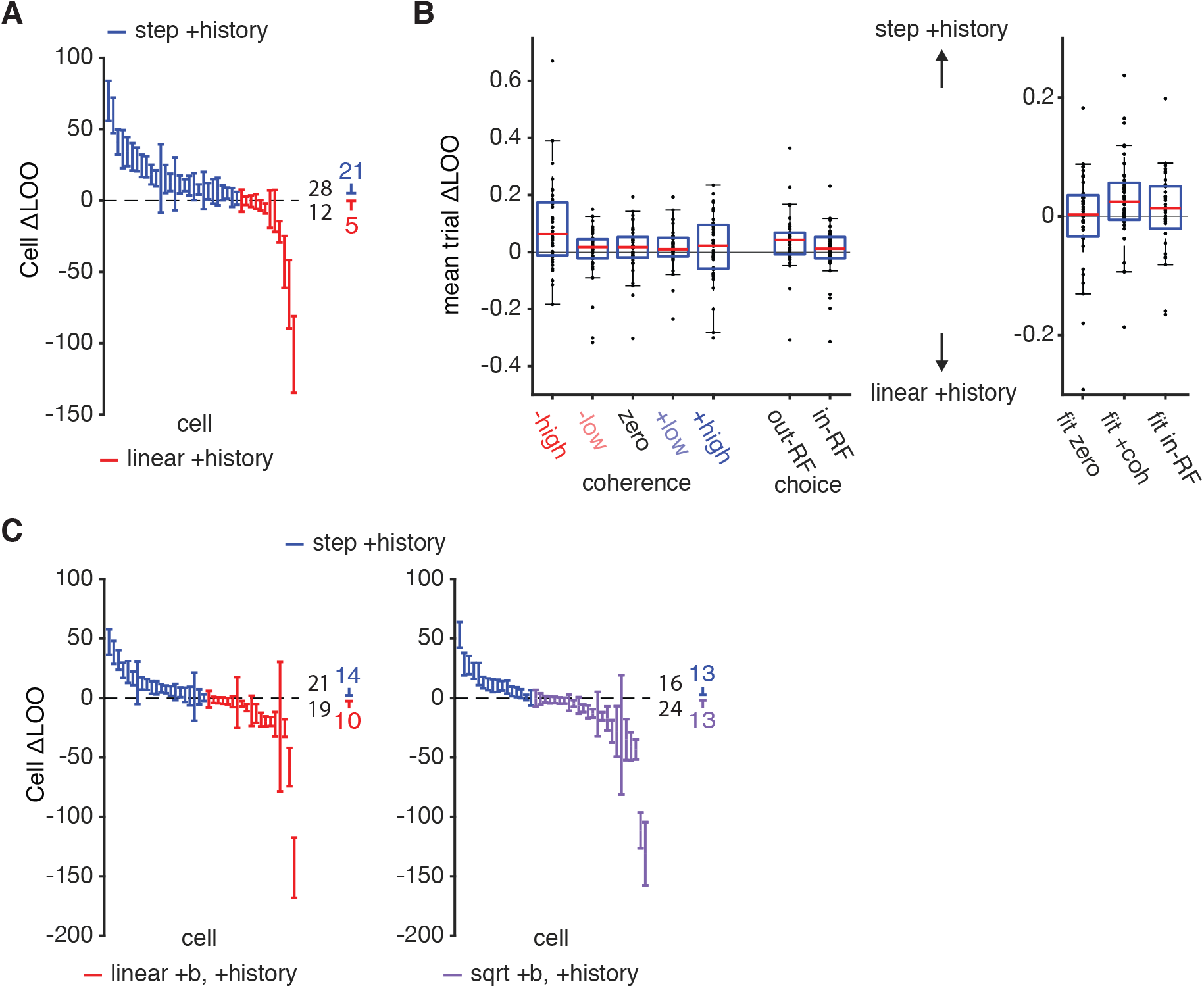
Model comparison using leave-one-out cross validation. Related to Figures 3 and 4. We used PSISLOO to estimate the Bayesian leave-one-out cross validation performance (LOO) on held out trials (Vehtari et al., 2017). In Bayesian LOO, predictive performance on a held-out trial is evaluated by integrating over the posterior distribution of the parameters conditioned on the rest of the trials. PSIS-LOO uses importance sampling to estimate this intergral. The proposal distribution is the posterior distribution conditioned on all of the trials (which we sample from in our MCMC procedure) and the importance weights are stabilized through regularization. Model comparison using LOO provided nearly identical results to the model comparison using WAIC.

**Figure 9.**
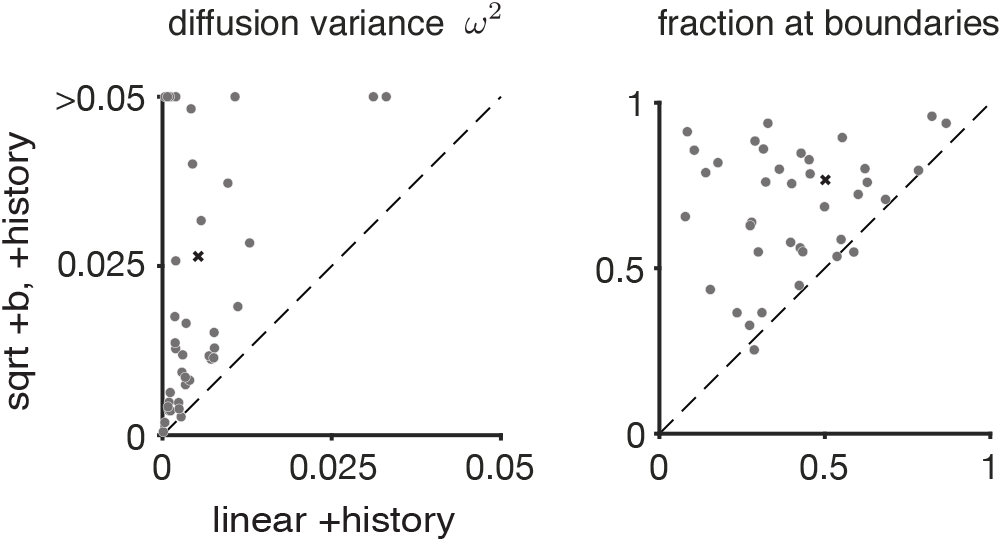
Related to Figure 5. Quantitative comparison of simulated trajectories from the square root ramping model with non-zero baseline and history against the linear ramping model with history and zero baseline. *Left:* Diffusion variance increased in the square root ramping model with non-zero baseline. *Right:* Fraction of time the simulated latent firing rates were equal to the baseline rate or upper absorbing boundary. Cell 1 is marked by an (x) in these plots.

**Figure 10.**
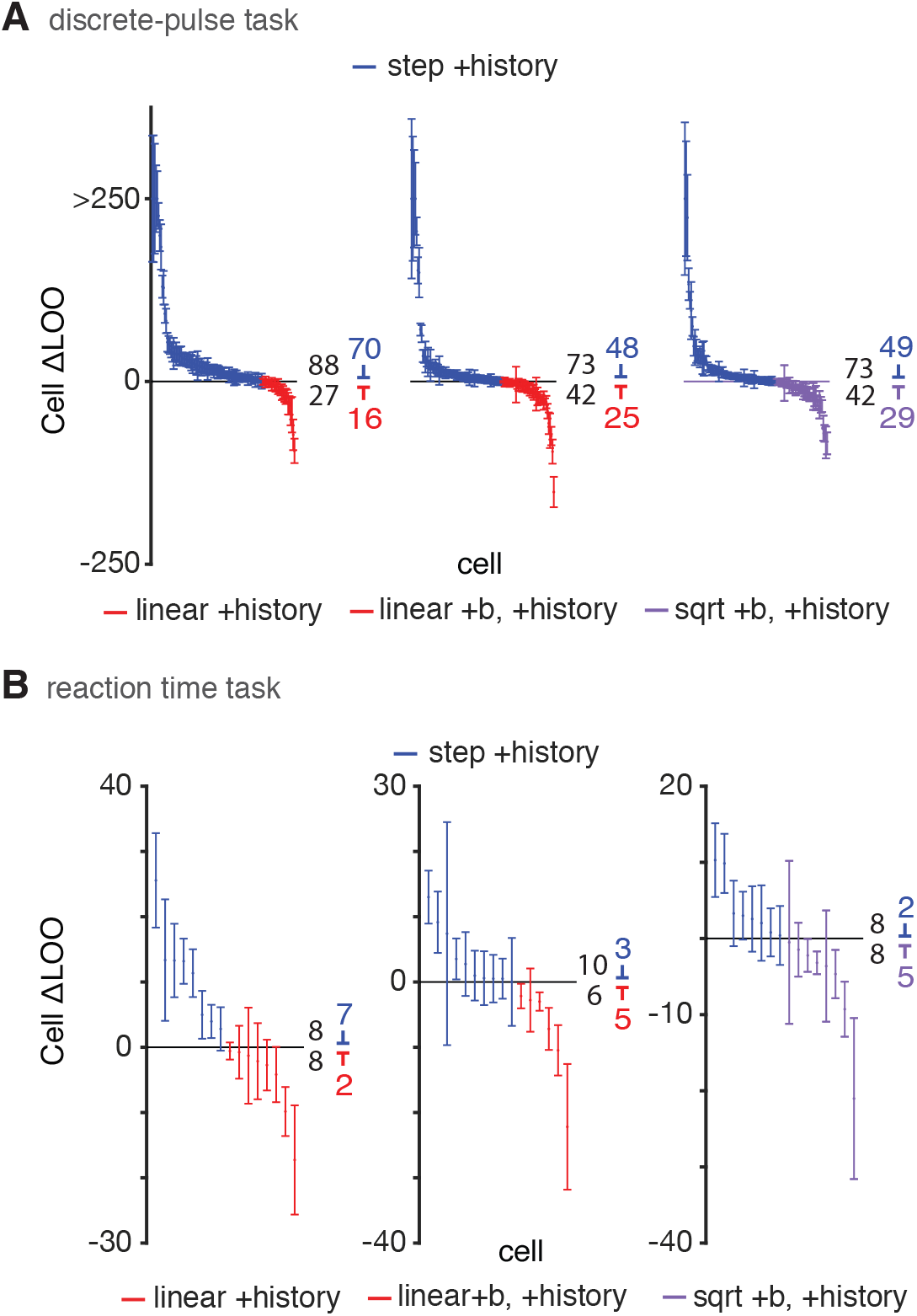
Model comparison using leave-one-out cross validation (LOO) for the discrete-pulse and reaction time data. Related to Figure 6. LOO was estimated using PSIS-LOO (Vehtari et al., 2017, see Figure 8). Model comparison using LOO for these datasets provided very similar results to model comparison using WAIC.

